# Drought induced metabolomics of potato leaves highlight metabolic reprogramming and promising biomarkers for smart irrigation advisories

**DOI:** 10.64898/2026.03.19.712810

**Authors:** Portia D Singh, Ramnath Nayak, Yvonne Dittrich, Radoslaw Guzinski, Yogesh Pant, Shyam Kumar Masakapalli

## Abstract

Smart irrigation management is essential for improving crop resilience under increasing drought frequency driven by climate change. Although satellite-based remote sensing provides valuable tools for monitoring crop water status at large spatial scales, its accuracy is often limited in mountainous and heterogeneous agricultural landscapes. In this study, we investigated drought-induced metabolic responses in potato (*Solanum tuberosum* L.) to identify biochemical biomarkers that could complement satellite-based irrigation advisories in the mid-Himalayan region of India. A field experiment was conducted using a gradient of soil moisture regimes corresponding to moderate (50% field capacity), critical (25% field capacity), and extreme drought stress (5-8% field capacity). Satellite-derived evapotranspiration-based irrigation advisories were validated against in situ soil moisture measurements, revealing discrepancies attributed to the inability of satellite estimates to capture actual water loss under drought stress conditions, highlighting the need for additional ground-truth biomarkers across heterogeneous field conditions. To capture plant-level physiological responses, untargeted metabolite profiling of potato leaves was performed using gas chromatography–mass spectrometry (GC-MS). Approximately fifty metabolites belonging to amino acids, organic acids, sugars, and sugar alcohols were detected. Multivariate statistical analyses revealed distinct metabolic signatures associated with progressive drought stress. Notably, accumulation of proline, serine, isoleucine, sucrose, fructose, glucose, and polyols such as mannitol and myo-inositol reflected key metabolic reprogramming associated with osmoprotection, redox homeostasis, and energy metabolism under drought conditions. Collectively, this ensemble of stress-responsive metabolites represents a robust panel of drought stress biomarkers. As a proof of concept, proline was validated as a qualitative biomarker of plant water status through a rapid and cost-effective colorimetric biochemical assay, demonstrating its practical applicability for field-level irrigation management. These findings demonstrate that metabolomics-derived biomarkers can provide sensitive plant-level indicators of drought stress that complement satellite-based monitoring systems. The integration of biochemical diagnostics with remote sensing platforms offers a promising approach for improving drought detection and developing low-cost, field-deployable tools for smart irrigation advisories in heterogeneous agricultural landscapes.

**Graphical abstract:** 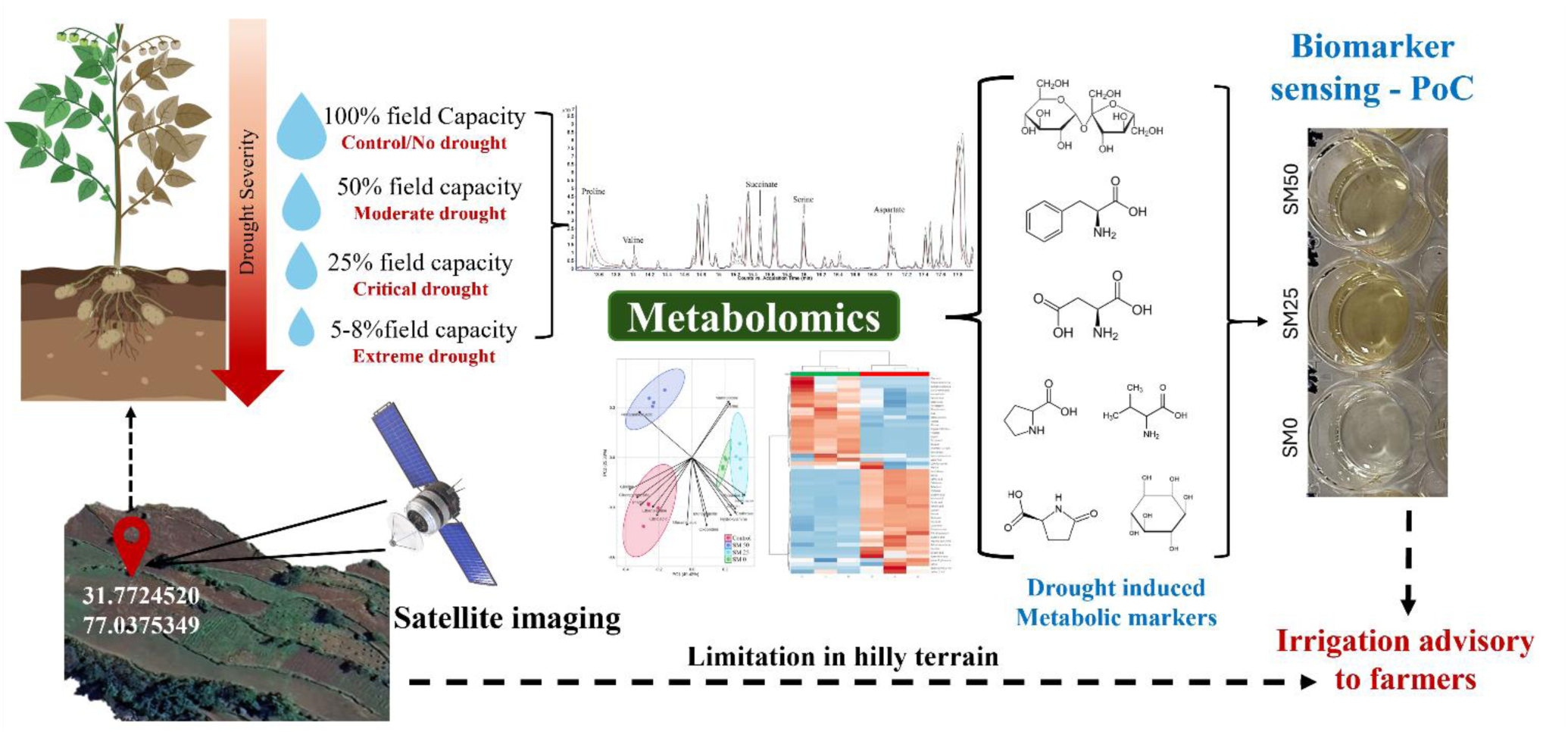

## 1 Introduction

Potato (*Solanum tuberosum* L.) ranks among the most important staple food crops globally, providing essential calories and nutrients to billions. However, its production is significantly vulnerable to abiotic stresses, particularly drought, which severely limits tuber yield and quality (Obidiegwu et al., 2015). The increasing frequency and severity of drought events due to climate change necessitate the development of advanced, efficient methods for early stress detection to enable timely irrigation management and mitigate crop losses (Satapathy et al., 2024; Disasa et al., 2025). Traditional approaches for assessing water stress often rely on visual symptoms, which typically manifest only after irreversible physiological damage and yield potential reduction have occurred. While physiological measurements offer more sensitivity, they can be labor-intensive and impractical for large-scale or rapid assessments (Jones; 2007). The era of Smart Agriculture offers powerful alternatives by leveraging data-driven technologies for precision crop management. Among these, satellite-based remote sensing provides invaluable tools for monitoring crop conditions across extensive agricultural areas. Techniques utilizing multispectral, hyperspectral, or thermal imagery can assess parameters like vegetation indices (e.g., NDVI), canopy temperature and evapotranspiration, which correlate with plant water status and photosynthetic activity, offering a macro-scale perspective on potential stress development (Ahmad et al., 2021). Although satellite remote sensing provides valuable spatio-temporal coverage for monitoring vegetation dynamics, its reliance on broad spectral indices limits the accurate identification of crop-specific stress responses. Because these signals are indirect and influenced by multiple environmental and biological factors, molecular or ground-based physiological measurements are necessary to validate and accurately interpret stress status (Wen et al., 2021 and Cooper et al., 2025). Metabolomics offers a multivariate understanding of plant biochemical status and, when integrated with satellite-based or high-throughput remote sensing, can enhance early detection and diagnosis of crop diseases by linking molecular responses to observable stress patterns (Arbona et al., 2013; Terentev & Dolzhenko, 2023).Metabolic read-outs offer systems-level insights into plant physiological adjustments to environmental conditions. Plants exhibit diverse phenotypic plasticity under water deficit, marked by early sensing and signaling, hormonal activation (primarily ABA), physiological and metabolic adjustments, structural adaptations, and eventual recovery or senescence depending on stress duration. Under drought stress, potato plants accumulate or deplete specific metabolites such as proline, soluble sugars, abscisic acid, and various amino acids as part of their adaptive response (Gong et al., 2015; Vasquez-Robinet et al., 2008). These molecules serve as sensitive biomarkers, enabling early and specific detection of water deficit at the cellular level, often before physiological changes become reliably detectable from space (Arbona et al., 2013; Obata & Fernie, 2012). Integrating the broad spatio-temporal monitoring capabilities of satellite-based imaging with the high sensitivity of metabolomic biomarkers provides a powerful, complementary approach for managing drought stress in potato. By linking field-scale remote sensing with precise biochemical indicators, this strategy enables more accurate, timely, and targeted interventions to enhance crop resilience and productivity under water-limited conditions (Ahmad et al., 2021; Obata & Fernie, 2012).

We hypothesize that metabolite-level responses in potato leaves provide early and sensitive biomarkers of drought stress that can complement satellite-based irrigation advisories in heterogeneous mountainous terrains. Therefore, in this study, we aimed to (i) demonstrate that satellite-based drought advisories require complementary support from high-resolution ground observations, particularly in hilly terrain with small and heterogenous agricultural fields, and (ii) establish metabolic biomarkers of drought stress in potato (*Kufri Jyoti*) by integrating soil moisture gradients with GC-MS-based untargeted metabolite profiling. These findings underscore the potential for developing low-cost, field-deployable biochemical assays for real-time drought diagnostics, thereby providing a complementary platform to remote sensing in topographically complex terrains and enabling more precise, data-driven irrigation advisories.

## 2 Material and Method

### 2.1 Plant material

The study was conducted in March- April 2024 during the potato (*Solanum tuberosum* cv. *Kufri Jyoti*) growing season in a field located in Sakaryar Village, Mandi, Himachal Pradesh, India (Latitude: 31.7724520, Longitude: 77.0375349). Two experimental plots were systematically prepared and fertilized to ensure adequate nutrient availability. Environmental data collection included daily recordings of minimum and maximum temperatures (°C) and average relative humidity (%) using a Tempnote TH32 Portable Temperature Data Logger. Soil moisture (%) was measured daily using a Lutron PMS-714 Soil Moisture Meter. The soil moisture readings were calibrated and correlated with field capacity (Supplementary Table 1). To complement ground-based observations, daily evapotranspiration (ET) data for the specified field coordinates were derived through satellite image analysis (Guzinski et al., 2020 & Guzinski et al., 2021), with the objective of establishing correlations between satellite-derived and field-measured environmental variables for deriving irrigation advisories (Supplementary Table 1).

### 2.2 Experimental design for water stress assessment

The water stress experiment was conducted over two weeks in the field (March 22 - April 5, 2024) at two locations: Plot A (control, regularly irrigated) and Plot B (subjected to drought stress). Drought conditions in Plot B were categorized into three soil moisture (SM) regimes based on field capacity (FC), calculated from daily weight loss: (1) moderate stress (SM 50, 50% FC), (2) critical stress (SM 25, 25% FC), and (3) extreme stress (SM 0, 5-8% FC).Satellite image analysis was performed using the Sen-ET modelling approach (Guzinski et al., 2020 & Guzinski et al., 2021), covering the experimental site. In this approach thermal observations from Sentinel-3 satellites (daily overpass, 1 km spatial resolution) are used synergistically with shortwave optical observations from Sentinel-2 satellites (overpass every 5 days, 20 m spatial resolution), meteorological forcing and landcover map to drive a physically based actual evapotranspiration (ET) model. The land cover map, used for ET model parameterization, was adjusted to reflect the location of the potato fields to enhance the accuracy of evapotranspiration and water deficit modelling. The output of the ET model were actual and potential evapotranspiration maps (in mm/day) with 20 m by 20 m pixel size. The maps were produced daily, however on days in which the potato field was obscured by clouds during the satellite data acquisition, a gap-filling approach was used instead of the physical model (Guzinski et al., 2021). The gap-filling uses the last available clear-sky ET estimates and accounts for changing meteorological conditions (including solar irradiance) but does not consider soil drying or wetting during. To estimate water stress, water deficit was calculated, defined as (potential ET – actual ET).

### 2.3 Proline content test

Proline quantification in potato leaf samples was performed using the RealGene™ Proline Content Estimation Kit, following the manufacturer’s recommended protocol. Approximately 100 mg of fresh leaf tissue was homogenized in 1 mL of extraction reagent and centrifuged at 10,000 × g for 10 minutes at 4°C. From the resulting supernatant, 500 µL was combined with 500 µL of Reagent 1 (glacial acetic acid) and 500 µL of Reagent 2, followed by incubation at 100°C for 60 minutes. After vortexing, the mixture was subjected to spectrophotometric analysis, with absorbance measured at 520 nm. Proline concentration was determined using a standard calibration curve generated from L-proline standards, and results were expressed as micromoles per gram of fresh weight (µmol g⁻¹ FW). All measurements were conducted in biological replicates (n = 4) to ensure statistical reliability. For comparison, L-proline standard solutions ranging from 0 to 40 µg/mL were prepared.

### 2.4 Metabolite Profiling Workflow: Sample Preparation, Metabolite Extraction, and GC-MS Data Acquisition

To profile metabolites, lyophilized potato leaf tissue (∼50 mg) was used for soluble metabolite extraction following the protocol described by Lisec et al., (2006). Each sample was homogenized in 940 µL of extraction solvent comprising methanol:chloroform:water (3:1:1, v/v/v) and supplemented with 60 µL of ribitol (0.2 mg/mL in H₂O) as an internal standard. Samples were incubated at 70°C for 5 minutes in a thermomixer at 950 rpm, followed by centrifugation at 13,000 × g for 10 minutes at room temperature. Approximately 50 µL of the supernatant was transferred to a fresh microcentrifuge tube and vacuum dried. Dried extracts were derivatized via methoxyamine hydrochloride (MeOX) and trimethylsilyl (TMS) reagent treatment, as described by Masakapalli et al., (2014). Samples were subsequently analyzed using a GC-MS system (GC ALS-MS 5977B, Agilent Technologies) equipped with an HP-5ms capillary column (30 m × 250 μm × 0.25 μm). Metabolite identification was based on comparison of mass ion fragments (m/z), retention times (RT), and characteristic identifier ions with spectral data from the NIST 17 library. For statistical and multivariate data analysis, we utilized MetaboAnalyst 6.0, accessible at https://www.metaboanalyst.ca/. Data pre-processing was performed by normalization by the median, log transformation, and Pareto scaling. The area of the internal standard, Ribitol, is used to obtain the relative proportions of the peaks, leading to the calculation of fold changes of metabolite/ peak levels among the treatments.

## 3 Results and Discussion

### 3.1 Integration of satellite image analysis with ground-truth measurements on potato field crop for enhanced drought monitoring in mountainous terrain

Daily environmental parameters for field experiments were recorded, including minimum and maximum temperatures (°C), average relative humidity (%), and soil moisture content (%) (Supplementary Table 2). Notably, discrepancies emerged between ground-based soil moisture data and satellite-derived evapotranspiration data. While satellite-based advisories indicated “watered field” conditions, ground measurements revealed a progressive decline in soil moisture to 50-25% of field capacity (Figure 1, Supplementary Table 1). Satellite advisories continued signaling adequate moisture across drought-stressed plot until 2 April 2024, failing to detect localized water stress under both mild (SM 50) and critical (SM 25) drought conditions. Only after this date did satellite observations indicate irrigation requirements (Supplementary Table 3). There are several potential sources for this inconsistency. Satellite image analysis in mountainous regions like Himachal Pradesh is challenging due to complex topography, variable land cover, frequent cloud cover, and small fragmented plots compromising remotely sensed data accuracy (Ma et al., 2025; Chen et al., 2023; Zhu et al., 2013; Zhu et al., 2012). Specifically, during the experiment there was a gap in cloud-free satellite observations between 25/03/2024 and 02/04/2024. Therefore, the transition from mild drought to extreme drought for plot B (Figure 1) was not observed by satellites and instead the gap-filled ET values, which do not consider soil drying, were used for satellite-based irrigation advisory. As soon as satellite data become available, the advisory changed to “Required

**Figure 1:**
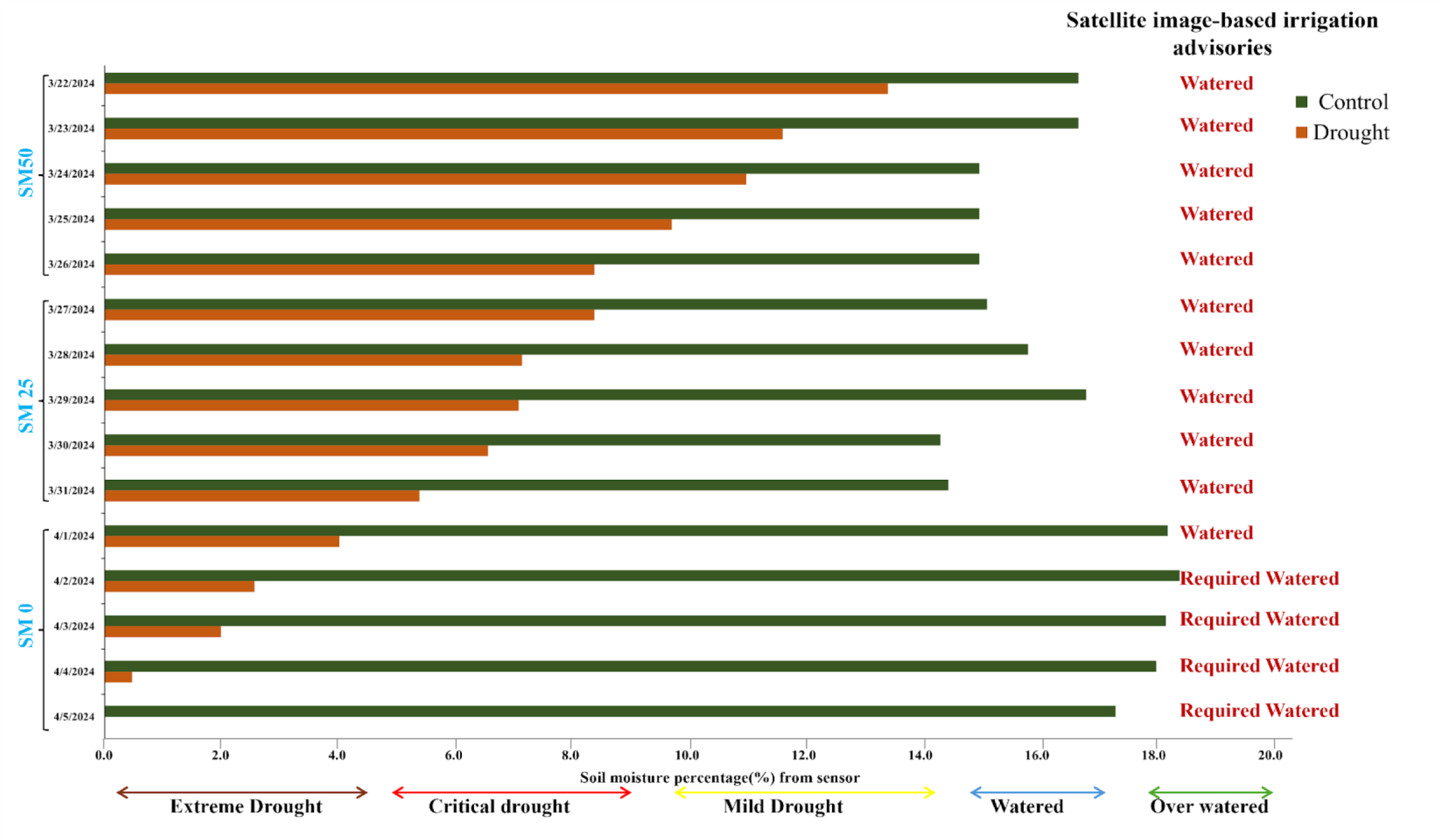
Validation of satellite data and actual soil moisture level under different drought regimes. Soil moisture reading from sensors and ET data advisories from March 22, 2024, to April 5, 2024.The green bar showed the soil moisture value at control field, orange bars represent soil moisture values in drought field, the red text (watered /required water) represent the irrigation advisories based on ET values derived from satellite image analysis. Notably, the satellite-based advisories failed to detect localized water stress in drought fields until 2 April 2024, despite declining in-situ soil moisture levels (SM 50 and SM 25).

Watered”. Additionally, advisories derived from 20×20 m spatial resolution data represented mean values across multiple small fields within the same pixel. Sparse ground-truth data in such terrains further hinders proper calibration and validation of satellite models (Brocca et al., 2011). Finally, the analysis relied on a generic vegetation response model to translate modeled potential and actual evapotranspiration into plant water stress. However, as demonstrated by our results, potato being a tuber crop exhibits a crop-specific water response function that differs substantially from that of generic vegetation parameterizations. Consequently, satellite-based drought advisories in such agro-ecological settings should be complemented with high-resolution ground observations and crop-specific, plant-level diagnostics to account for physiological and phenological differences, thereby improving the reliability and accuracy of drought stress assessment.

### 3.2 Multivariate analysis reveals distinct metabolic fingerprints across progressive drought stress levels

GC-MS-based metabolite profiling of the potato leaves detected approximately 50 metabolites belonging to the classes of amino acids, fatty acids, organic acids, sugars, and sugar alcohols (Supplementary Table 4). Multivariate analyses revealed clear metabolic variations among control, and drought treatments of SM 50 (moderate stress), SM 25 (critical stress), and SM 0 (extreme stress) (Figure 2). PCA captured 74.75% of variance across PC1 and PC2, effectively separating the treatment groups (Figure 2.A). Control samples are characterized by higher glycine, glucopyranoside, pinitol, ethanolamine, and citric acid. Pentanedioic acid strongly associates with SM 50, while mannobiose and serine contribute to its separation. SM 25 and SM 0 exhibit elevated threonine, isoleucine, arbutin, erythrose, hydroxylamine, monopalmitin, oxoproline, and mannobiose, indicating stress-related metabolic shifts. The VIP score plot identified proline as the most discriminant metabolite, peaking under SM 25, suggesting a strong adaptive response to moderate drought (Figure 2.B). Changes in amino acids (proline, valine, tryptophan, serine, glycine, threonine) reflected osmoprotection, stress signaling, and biosynthetic roles in drought-stressed plants (Obata & Fernie, 2012). Proline accumulation, in particular, serves multifunctional roles in stress tolerance (Szabados & Savouré, 2010), while branched-chain and aromatic amino acids participate in broader metabolic reprogramming (Less & Galili, 2008). Variations in organic acids (citric, fumaric, succinic, citramalic) reflected TCA cycle adjustments and energy production under drought stress (Araújo et al., 2012; Nunes-Nesi et al., 2010), consistent with central carbon-nitrogen metabolic regulation in photosynthetic tissues. Pinitol variation tied to osmotic stress tolerance responses in potato (Chourasia et al.,2021), functioning as a compatible solute and osmoprotectant. PLS-DA biplot clearly separates groups, highlighting citric acid and pinitol in control, tryptophan in SM 50, proline and valine in SM 25, and phytol in SM 0 (Figure 2.C). Hierarchical clustering reveals tight within-group similarity with distinct treatment-based clustering (Figure 2.D). Notably, SM 25 and SM 0 cluster closely, as do control and SM 50, suggesting shared metabolic responses within pairs. These analyses confirm drought stress induced metabolic reprogramming, particularly affecting amino acid and organic acid pools reflective of adaptive physiological responses.

**Figure 2:**
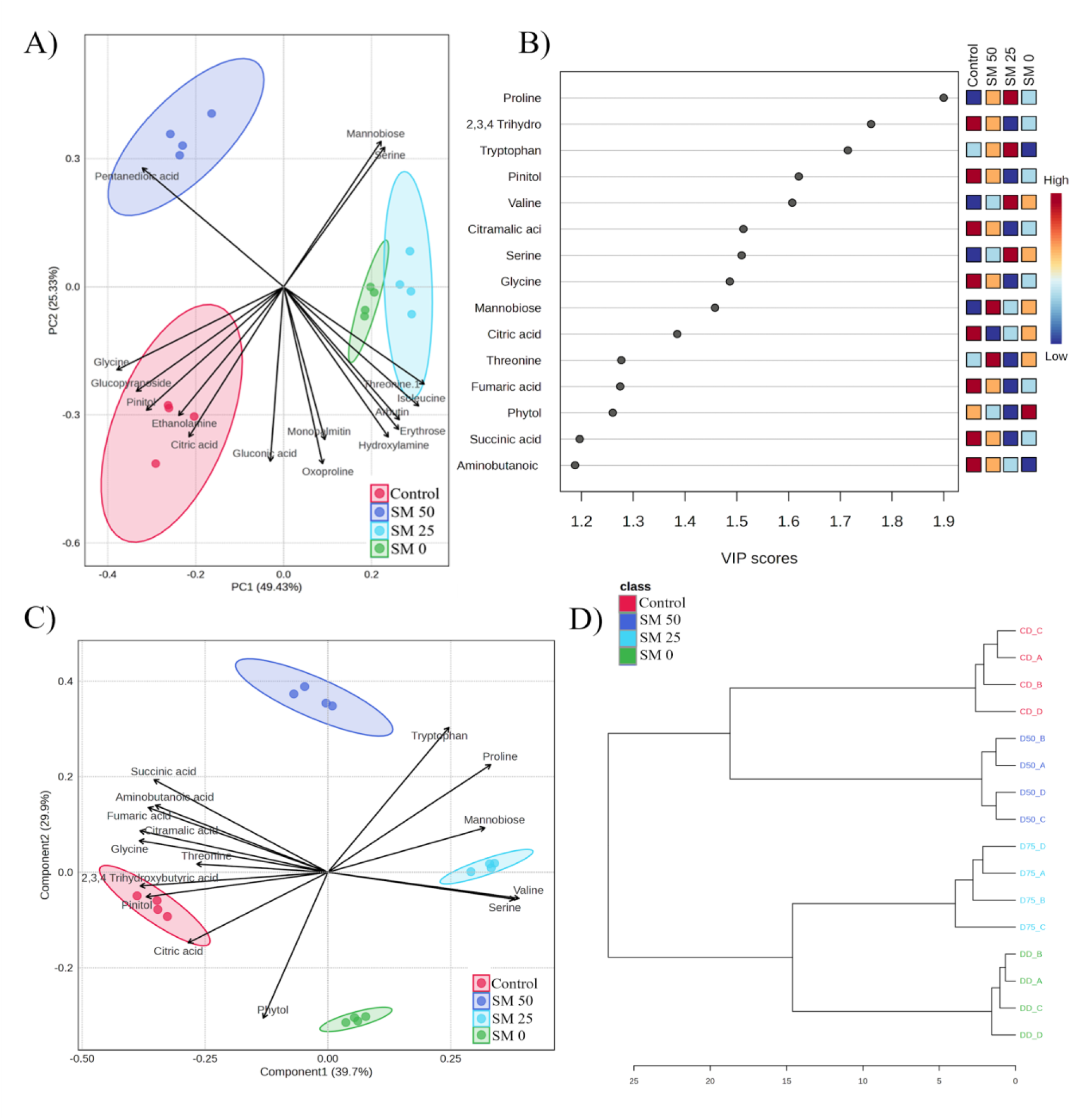
Metabolites captured in potato leaves under drought treatment showed varied metabolomes. (A) Principal component analysis (PCA) plot showing the separation of different treatments based on metabolite profiles. Each treatment group (Control, SM 50, SM 25, SM 0) is represented by an ellipsoid, and key metabolites contributing to the variation are labeled. (B) Variable Importance in Projection (VIP) scores of metabolites identified through partial least squares (PLS) analysis, ranked by their contribution to the class separation. Colors indicate the relative abundance of metabolites across different treatment groups. (C)The PLS-DA biplot for component 1 (39.7%) and component 2 (29.9%), showing the distribution of metabolites in relation to experimental treatments. Significant metabolites contributing to each group are highlighted aligning with the VIP analysis results. (D) Hierarchical clustering dendrogram based on metabolite profiles, showing the clustering of treatment groups. The dendrogram indicates similarities and differences in metabolic profiles between the control and treatment groups (SM 50, SM 25, SM 0).

### 3.3 Drought-induced amino acid accumulation in potato leaves is a key adaptive response to declining soil moisture

Progressive drought stress triggered substantial metabolic reprogramming in potato leaves, particularly marked by amino acid accumulation peaking under critical stress (SM 25), with significant increases in proline, valine, serine, threonine, tryptophan, and phenylalanine (Figure 3). Proline showed the most dramatic rise, functioning as a central osmoprotectant and stress marker (Szabados & Savouré, 2010; Singh et al., 2015). Amino acid accumulation served multiple adaptive functions: supporting osmotic adjustment by lowering cellular water potential to maintain turgor (Chaves et al., 2003), accumulating from drought-induced growth inhibition and enhanced protein degradation (Hildebrandt et al., 2015), and providing ROS scavenging and membrane stabilization. Aromatic amino acids like phenylalanine and tryptophan serve as precursors for defense compounds via the shikimate pathway (Galili et al., 2016; Tzin & Galili, 2010), while accumulated amino acids may represent temporary nitrogen storage under stress conditions (Less & Galili, 2008). Distinct response patterns including early aspartic acid peak at SM 50, continued isoleucine increase to SM 0, and glycine decrease highlighted the complexity of amino acid metabolic regulation in potato leaves under progressive drought stress. These differential accumulation patterns likely reflect amino acid-specific pathway regulation, differential stress signal transduction, and dynamic shifts in metabolic flux priorities as drought stress intensified (Galili et al., 2016; Less & Galili, 2008). Such coordinated but differentiated amino acid responses represent a critical adaptive strategy in potato, integrating osmotic regulation, resource management, cellular protection, and signaling pathways to enhance plant survival during water scarcity.

**Figure 3:**
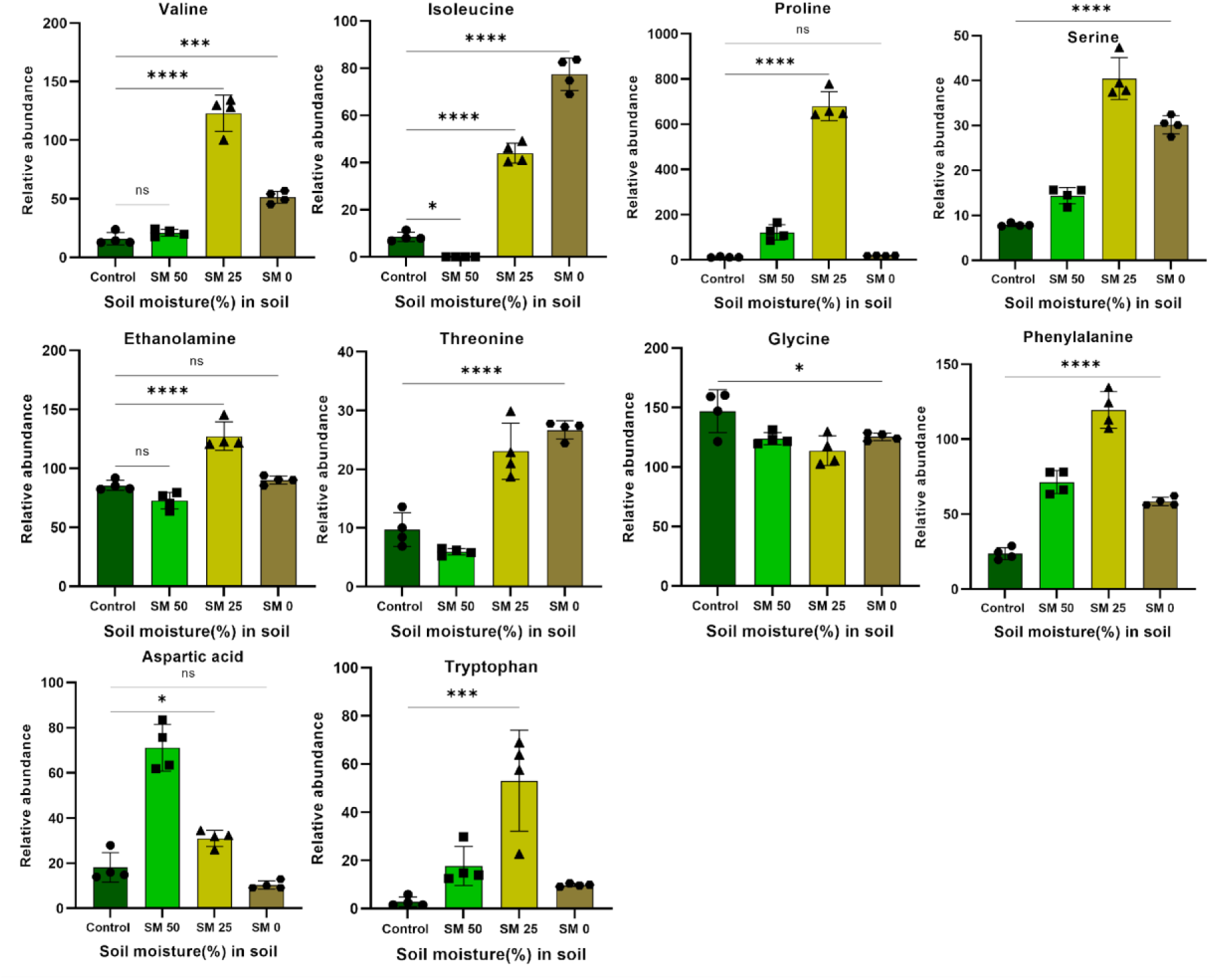
Drought exposure modulates amino acid metabolism in potato leaves. The levels of amino acids (aspartic acid, tryptophan and phenylalanine) are altered during SM 50 treatment. Higher increment of other amino acids was observed under SM 25 treatment. Data are presented as mean ± standard error mean (n = 4). Significant differences between groups were determined by statistical analysis: ns = not significant, *p < 0.05, **p < 0.01, ***p < 0.001, ****p < 0.0001.

### 3.4 Alterations within nitrogen metabolism and its pathways to varying drought stress intensities

Under drought stress in potato leaves, oxoproline accumulated significantly under critical-to-extreme conditions (SM 25 & SM 0) (Supplementary Figure S1.A), likely reflecting increased proline synthesis and cycling for osmoprotection, consistent with proline-glutamate pathway responses characterized across plant species (Verbruggen & Hermans, 2008; Szabados & Savouré, 2010). Notably, GABA levels remained stable across all drought severities, despite GABA being commonly induced by abiotic stress (Ramesh et al., 2017). This unexpected stability in potato leaves may indicate high metabolic turnover via the GABA shunt (Michaeli & Fromm, 2015) or reflect potato-specific regulation of nitrogen metabolism under water deficit. Hydroxylamine displayed a biphasic response, declining sharply under mild stress (SM 50) then spiking at critical stress (SM 25), indicating complex, stress-intensity- dependent shifts in nitrogen metabolism. As hydroxylamine participates in nitric oxide metabolism and nitrogen redox pathways (Astier et al., 2018; Gupta et al., 2011; Mur et al., 2013), its dynamic accumulation pattern in potato leaves likely reflects disrupted nitrogen flux or altered enzymatic activity under progressive drought conditions. These nitrogen metabolites collectively highlight specific, stress-intensity-dependent metabolic reprogramming in potato leaves responding to water scarcity.

### 3.5 Alterations in fatty acids metabolic pathways of potato leaves in response to drought stress

Significant alterations in fatty acid metabolism and signaling pathways occurred in potato leaves in response to declining soil moisture. Saturated fatty acids (palmitic, stearic) and their monoacylglycerols (monopalmitin, glycerol monostearate) decreased under moderate stress (SM 50) but peaked under critical-to-extreme stress (SM 25,SM 0) (Supplementary Figure S1.B). This accumulation likely reflected drought-induced membrane lipid remodeling, with increased saturated fatty acid proportions decreasing membrane fluidity, an adaptation commonly observed under abiotic stress in leaf tissues (Upchurch, 2008; Gigon et al., 2004). Conversely, the polyunsaturated fatty acid linolenic acid (C18:3) substantially decreased under all drought levels (SM 50, SM 25, SM 0), potentially due to increased lipid peroxidation from oxidative stress (Noctor et al., 2014), selective degradation of chloroplast membranes where linolenic acid is highly enriched, or diversion into jasmonic acid biosynthesis (Wasternack & Hause, 2013). Additionally, progressive accumulation of the biogenic amine octopamine under critical (SM 25) and extreme (SM 0) drought conditions suggested activation of specific stress signaling cascades in potato leaves. Catecholamines including octopamine and tyramine modulate stress responses in potato through effects on metabolism and physiology (Swiedrych et al., 2004; Kulma & Szopa, 2007), potentially influencing the observed metabolic adjustments under water deficit. Together, these coordinated changes in saturated lipids, unsaturated fatty acid depletion, and signaling amine accumulation highlight the intricate biochemical adjustments in lipid metabolism and stress signaling employed by potato leaves to cope with water deficit.

### 3.6 Metabolic reprogramming of central carbon metabolism and sugar accumulation under progressive drought stress

Drought stress significantly perturbs central carbon metabolism, revealing nuanced metabolic reprogramming rather than uniform responses (Supplementary Figure S2). TCA cycle intermediates show varied patterns: significant reductions in succinic acid (SM 25, SM 0) and malic acid (SM 0) likely result from reduced respiratory demand (Atkin & Macherel, 2009), limited CO₂ uptake affecting acetyl-CoA supply (Lawlor & Cornic, 2002), and diversion of carbon skeletons such as oxaloacetate and α-ketoglutarate toward synthesis of osmoprotective amino acids (Less & Galili, 2008; Araújo et al., 2012; Hildebrandt et al., 2015).Citric acid decreased under mild stress (SM 50) but recovered under stronger stress, potentially reflecting its multifunctional roles in cytosolic biosynthesis, cellular pH homeostasis, and metabolic regulation (Igamberdiev & Eprintsev, 2016; Sweetlove et al., 2010). Fumaric acid and citramalic acid remained stable, suggesting tight metabolic regulation or lower sensitivity to drought stress.

Carbohydrate metabolism in potato leaves showed multifaceted shifts under drought stress (Figure 4). Under mild stress (SM 50), sucrose, glucose, and fructose accumulated significantly, consistent with source-sink imbalance where growth inhibition outpaced reduced photosynthetic carbon fixation (Lawlor & Cornic, 2002). As stress intensified (SM 25, SM 0), these sugars declined due to increased conversion into protective compounds including amino acids and polyols (Hare et al., 1998), elevated energy demands for cellular maintenance, and photosynthetic feedback inhibition (Paul & Foyer, 2001). Under moderate-to-severe stress, accumulation of myoinositol, mannobiose, and erythrose reflected active carbohydrate metabolic reprogramming. Myoinositol and related polyols function as compatible solutes in drought-stressed potato leaves (Legay et al., 2011), serving as precursors for osmoprotectants like pinitol and galactinol while contributing to membrane stabilization and signaling (Valluru & Van den Ende, 2011). Polyols including mannitol contribute to ROS scavenging and oxidative protection (Smirnoff & Cumbes, 1989; Hare et al., 1998), processes critical for maintaining photosynthetic capacity under water deficit (Obidiegwu et al., 2015). Overall, these coordinated shifts highlight the metabolic flexibility of potato leaves, implementing strategic carbohydrate-to-polyol conversion supporting osmoprotection, energy balance, and stress adaptation (Legay et al., 2011; Pinheiro & Chaves, 2011).

**Figure 4:**
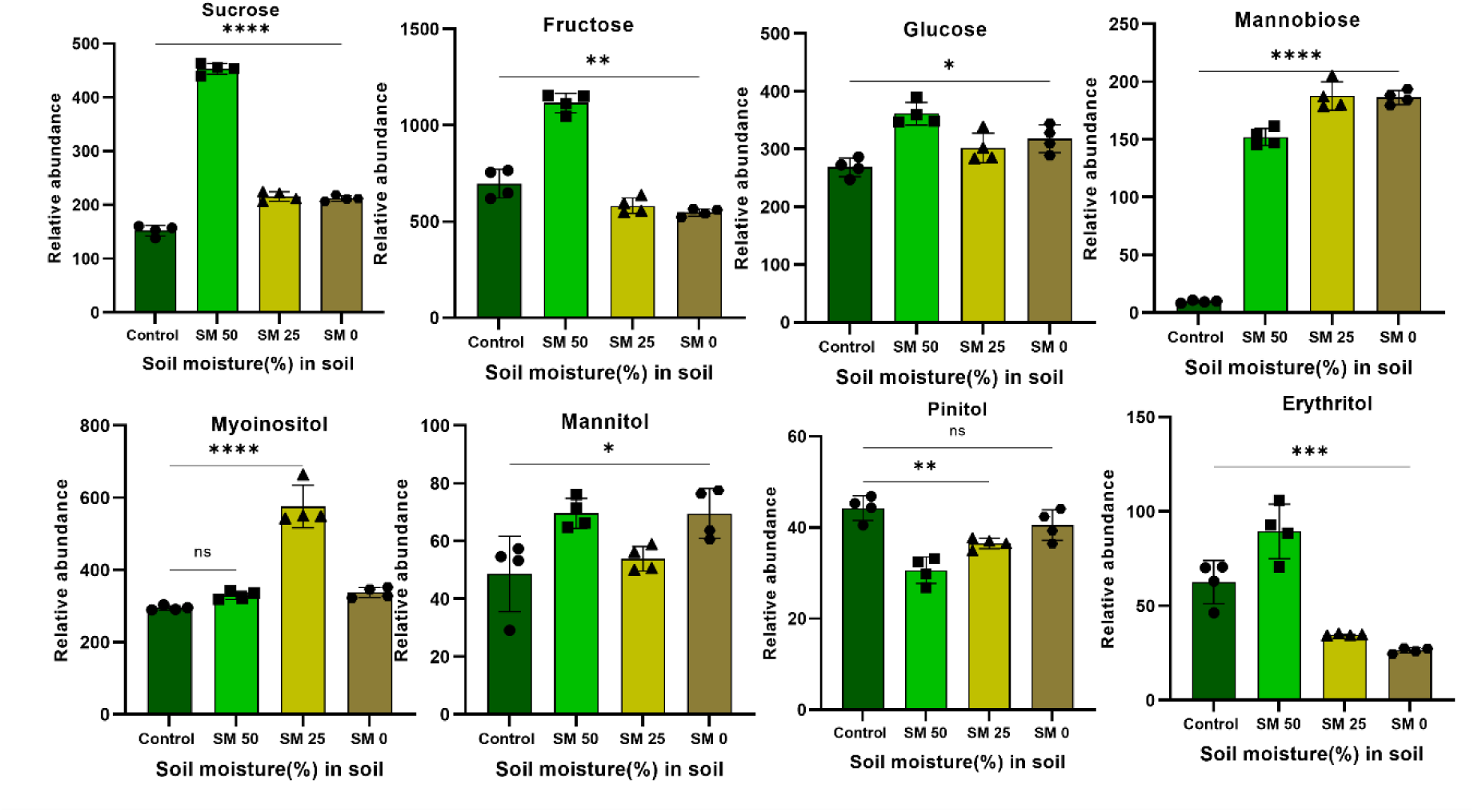
Kinetic drought exposure modulates sugar metabolism in potato leaves. Quantitative analysis of selected metabolites derived from GC-MS data across different soil moisture conditions: Control, SM 50, SM 25, and SM 0. The levels of soluble sugars (fructose, glucose, erythrose, sucrose, Mannitol, glucopyranoside) are altered in the mild stress (SM 50). Data are presented as mean ± standard error mean (n = 4). Significant differences between groups were determined by statistical analysis: ns = not significant, *p < 0.05, **p < 0.01, ***p < 0.001, ****p < 0.0001.

### 3.7 Drought stress induces metabolic shifts enhancing osmoprotection, energy balance, and stress resilience in plants

Drought stress induced extensive metabolic reprogramming in potato leaves to maintain cellular homeostasis and enhance stress tolerance (Figure 5). Primary affected pathways included glycolysis, photorespiration, the shikimate pathway, TCA cycle, GS/GOGAT cycle, branched-chain amino acid (BCAA) biosynthesis, and sugar metabolism, consistent with drought responses observed across potato genotypes (Sprenger et al., 2016; Vasquez-Robinet et al., 2008). Drought-induced accumulation of sucrose, glucose, fructose, pinitol, mannitol, and galactaric acid reflected altered osmoprotection and energy management (Evers et al., 2010; Legay et al., 2011). Elevated sucrose, mannitol, and erythritol levels indicated enhanced compatible solute accumulation to stabilize cellular structures and scavenge reactive oxygen species (Loescher, 1987; Hare et al., 1998). Photorespiration intensified under drought, evidenced by increased glycine-serine cycling, functioning in stress tolerance and redox balancing (Wingler et al., 2000). Shikimate pathway flux changes drove biosynthesis of phenylalanine, tryptophan, and phenolic compounds including arbutin, crucial for secondary metabolite production and defense signaling (Tzin & Galili, 2010; Sprenger et al., 2018). Enhanced BCAA and TCA-derived amino acid biosynthesis suggested prioritization of protein and osmolyte synthesis from altered TCA cycle carbohydrate flows (Hildebrandt et al., 2015; Araújo et al., 2012). Upregulated fatty acid biosynthesis (palmitic, stearic, monopalmitin, linolenic acids) reinforced membrane stability (Upchurch, 2008). The GS/GOGAT cycle showed increased glutamate, glutamine, and glutathione production for nitrogen assimilation and ROS protection (Less & Galili, 2008). TCA cycle intermediates (citrate, succinate, fumarate, malate, oxaloacetate) displayed varied accumulation patterns, reflecting adaptive changes in respiratory energy production and redox homeostasis (Atkin & Macherel, 2009; Sweetlove et al., 2010). Together, these metabolic shifts demonstrated a coordinated drought response in potato leaves, maintaining redox balance, cellular integrity, and adaptive capacity (Obidiegwu et al., 2015).

**Figure 5:**
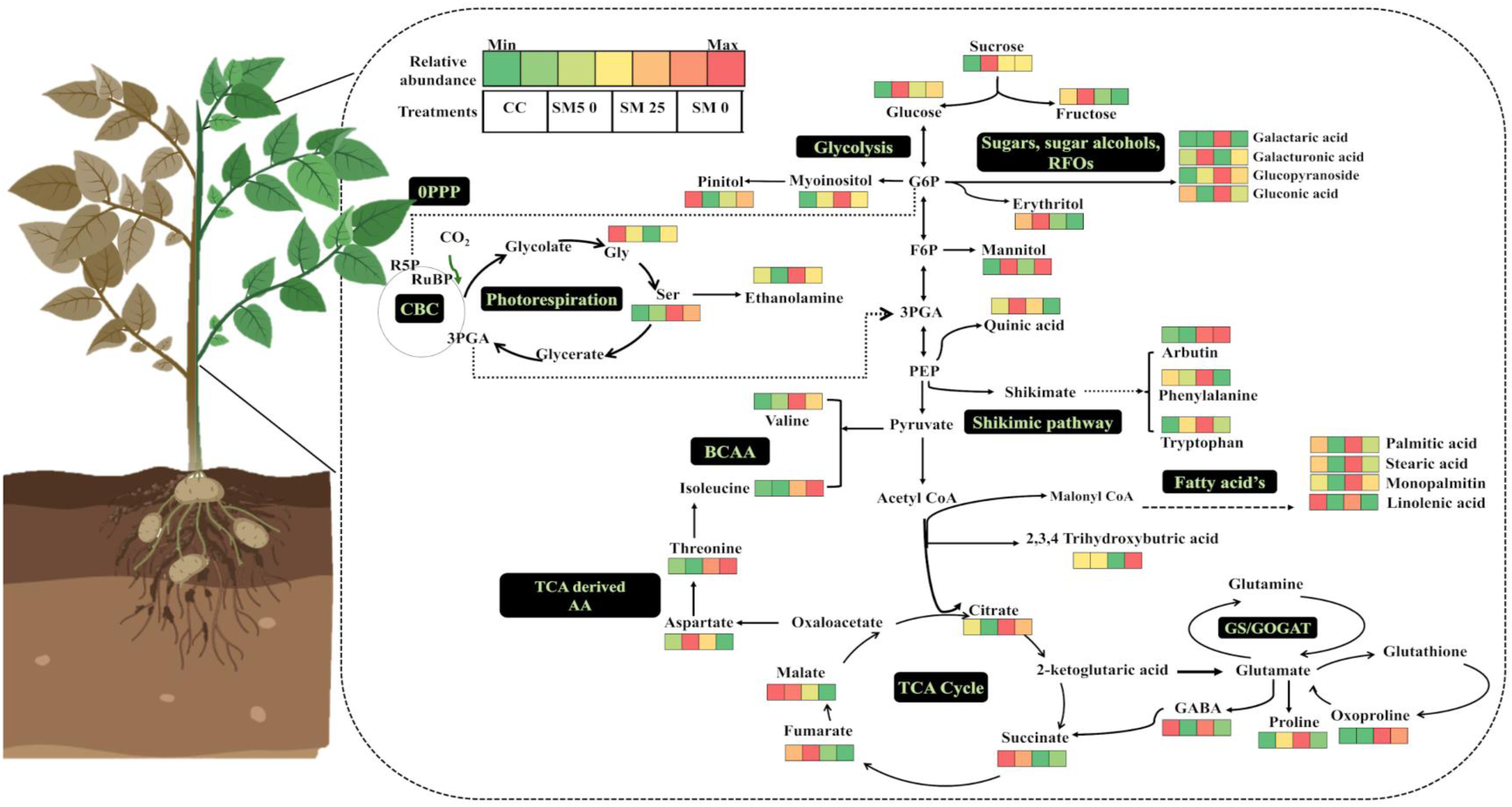
Drought induced metabolic variations in potato leaves. Relative abundance of metabolic change drought treatment was normalized with internal standard and average were means of four biological replicates. The altered pathways are shown as black colored boxes. The bar represents the Control (CC) and drought treatments (SM 50,25 and 0) metabolic response. The response of metabolites in the pathway is represented from Min to Max; dark red color has more abundance while green color is mapped to lower abundance. Altogether, few metabolites have similar responses to drought stress while levels of metabolites in GS/GOGAT, sugar and fatty acid metabolism were significantly changed.

### 3.8 Quantification of proline using an enzymatic and colorimetric assay kit

Gas chromatography-mass spectrometry (GC-MS)-based metabolic profiling is a powerful approach for elucidating plant responses to drought stress. However, GC-MS remains largely inaccessible to breeders and farmers unfamiliar with metabolomics. To bridge this gap, a simple, rapid, and cost-effective colorimetric assay for quantifying proline was evaluated to facilitate screening of drought-tolerant cultivars under field conditions (Figure 6.A). A clear physiological response to water deficit was observed (Figure 6.B), with proline accumulation increasing markedly as soil moisture decreased, consistent with its established role as a key osmoprotectant in plant drought tolerance. In field trials, proline content under critical drought stress (SM 25) was approximately two-fold higher than in well-watered controls (Figure 6.C). The distinct colorimetric changes associated with proline accumulation can be leveraged in two directions: (i) training machine learning models for integration into mobile-based diagnostic platforms, and (ii) developing low-cost, field-deployable biochemical assays for smart irrigation advisories (Figure 6.D). Notably, under extreme drought (SM0), proline levels remained comparable to controls, potentially indicating metabolic exhaustion or adaptive regulation where severe stress restricts further synthesis and accumulation (Bhaskar et al., 2015). While proline accumulation is not entirely drought-specific and may increase under other abiotic stresses such as salinity or extreme temperatures, and colorimetric assays can be influenced by interfering compounds, this assay remains a rapid and practical tool for field-level screening. These findings highlight proline’s reliability as a biochemical marker for drought stress and support its utility in screening drought-responsive cultivars (Szabados & Savouré, 2010; Singh et al., 2015). Furthermore, the broader ensemble of drought stress-responsive metabolites identified in this study shows a promising multi-biomarker platform that can be similarly employed for high-throughput physiological screening and precision drought stress assessment across diverse genotypes and field conditions.

**Figure 6:**
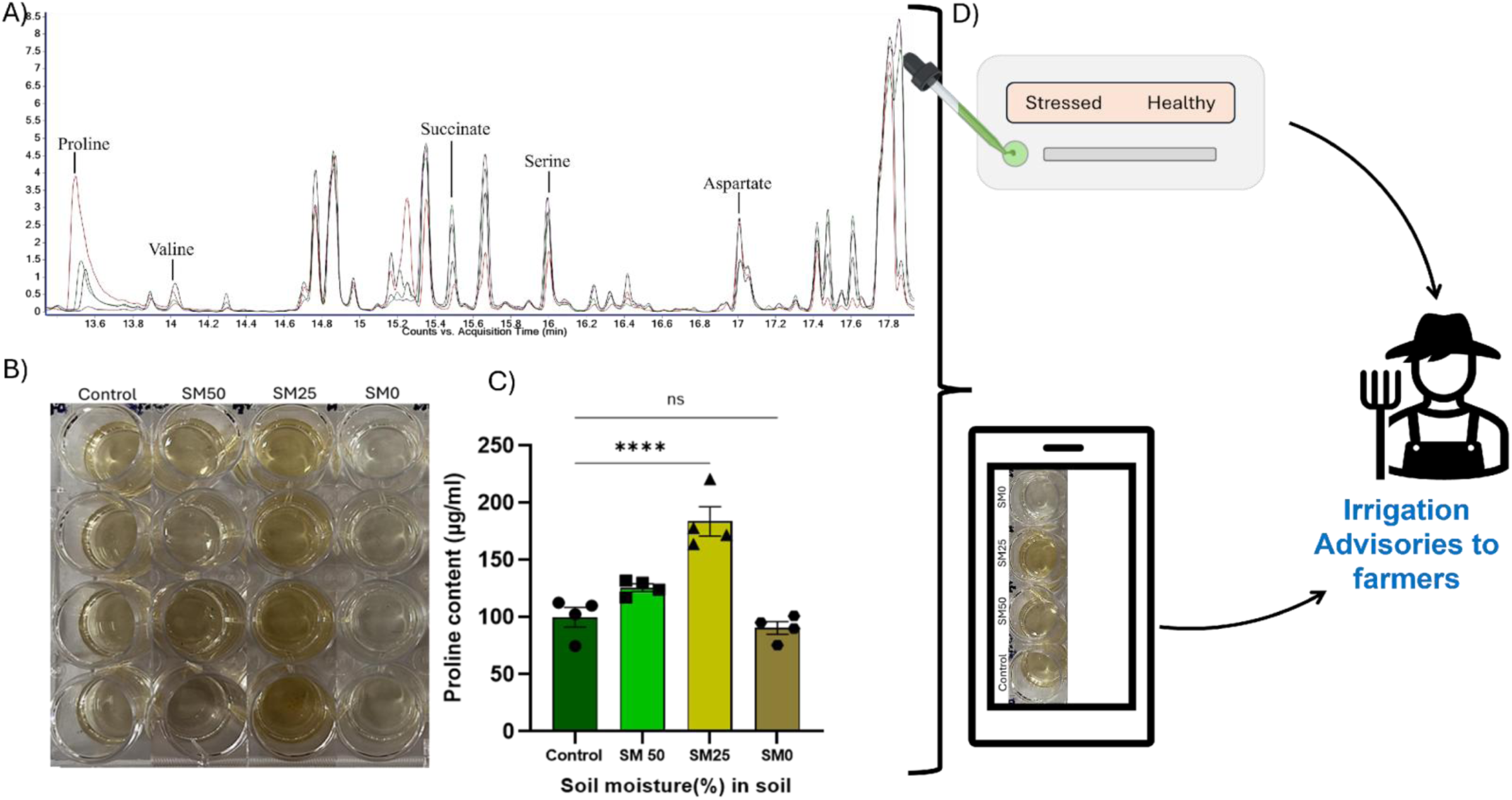
Drought biomarkers towards smart irrigation advisories, GC-MS, colorimetric, and biochemical based detection of drought stress for irrigation advisory. A) GC-MS spectra showing the relative abundance of metabolites Proline, Valine, Serine, and Aspartate under different soil moisture regimes, highlighting proline accumulation under drought stress. B) Colorimetric difference among the samples C) Proline content (µg/ml) in samples. Bars represent the mean values, with different symbols (circle, square, triangle, pentagon) indicating statistically significant differences between treatments (± standard error mean) **** =highly significant, ns = nonsignificant. D)A schematic representation of translation of assay adaptable for water stress diagnostics in future, aimed at supporting farmer-oriented irrigation advisory.

## 4 Conclusion

In conclusion, while satellite imagery provides valuable broad spatio-temporal coverage and performs reliably in plains, our findings reveal significant limitations in mountainous regions where complex topography, cloud cover, small and heterogenous agricultural fields and sparse calibration data hinder accurate drought detection. This study demonstrates that metabolomic-based biomarker analysis offers a robust complementary approach for drought stress management in potato crops, particularly in challenging terrains. Through GC-MS profiling, we identified a panel of early-stage (serine, isoleucine, pinitol) and late-stage (proline, myoinositol, valine, sucrose, fructose, glucose) metabolic biomarkers reflecting critical physiological processes such as osmotic adjustment and oxidative stress defence. Proline emerged as a particularly practical biomarker due to its consistent accumulation and simple colorimetric quantification. The proof-of-concept proline assay demonstrated field-level applicability through (i) integration with machine learning for mobile diagnostic platforms, and (ii) development of low-cost biochemical assays for timely irrigation advisories.

Overall, this study demonstrates the metabolomic biomarkers as a complementary strategy for drought detection, bridging plant physiology with practical agricultural tools. By enabling earlier and more precise monitoring in topographically complex regions where satellite advisories prove insufficient, these approaches can support optimized irrigation, improved water-use efficiency, and enhanced resilience of smallholder farming systems in water-scarce environments. Further, a better understanding of plant metabolic can be used to better calibrate satellite image analysis to the specific crop, even for plains.

## Acknowledgment

The authors are thankful to the SaIAFarm Team for all the data access. PDS acknowledge Mr Jagdish Chand (field staff) for continuous monitoring of experimental field, SKM acknowledge DST for funding support under INT_Denmark_P-4_2020 (G). PDS acknowledges project and PhD fellowships from the Ministry of Education and DST, Govt of India

## Contributions

PDS - conceptualization, data curation, formal analysis, validation, visualization, writing-original draft, RN - methodology, writing-review and editing, YD - funding acquisition, writing-review and editing, RG - satellite image analysis, writing- review and editing, YP-methodology, writing-review and editing, SKM - conceptualization, funding acquisition, supervision, writing-original draft, review and editing and PDS and SKM agree to be accountable for all aspects of the work.

## Declarations of interest

### Ethical approval

The study complied with the ethical standards.

### Data availability

The datasets generated during and/or analysed during the current study are available from the corresponding author on reasonable request.

