## Supplemental figures and tables for "Drought induced metabolomics of potato leaves highlight metabolic reprogramming and promising biomarkers for smart irrigation advisories"

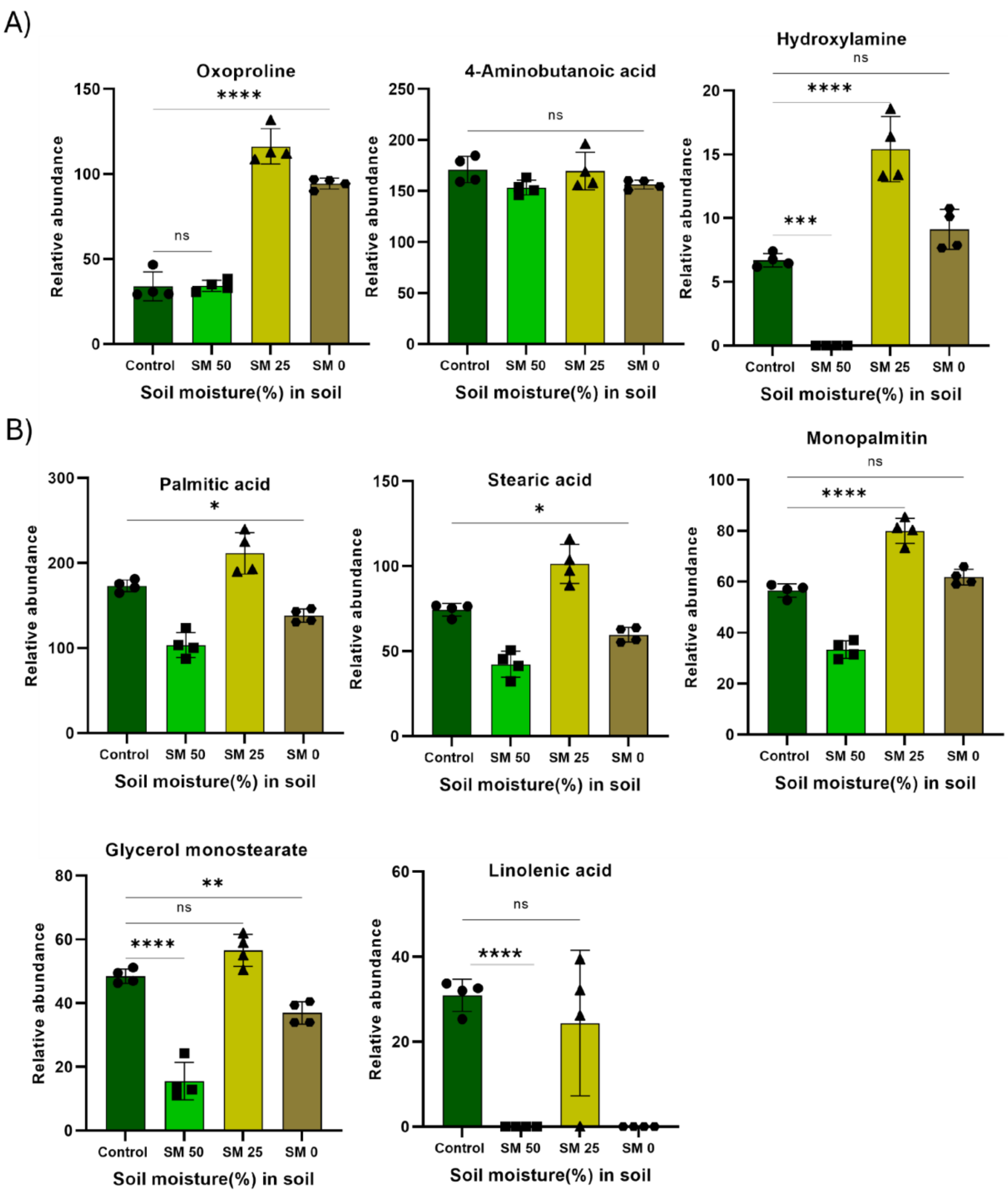


**Supplementary Figure S.1: Differential modulation of nitrogen and fatty acids metabolism in potato leaves under varying soil moisture levels.** A) Nitrogen metabolism compounds respond variably to soil moisture: Oxoproline and Hydroxylamine significantly increase at SM 25, while 4-Aminobutanoic acid remains unchanged. B) Lipid metabolism in potato leaves shifts with drought stress: palmitic and stearic acids, along with their monoacylglycerols (monopalmitin, glycerol monostearate), decline under mild stress (SM 50) but peak under moderate stress (SM 25) Data are presented as mean ± standard error mean (n = 4). Significant differences between groups were determined by statistical analysis: ns = not significant, *$p < 0.05$, **$p < 0.01$, ***$p < 0.001$, ****$p < 0.0001$.

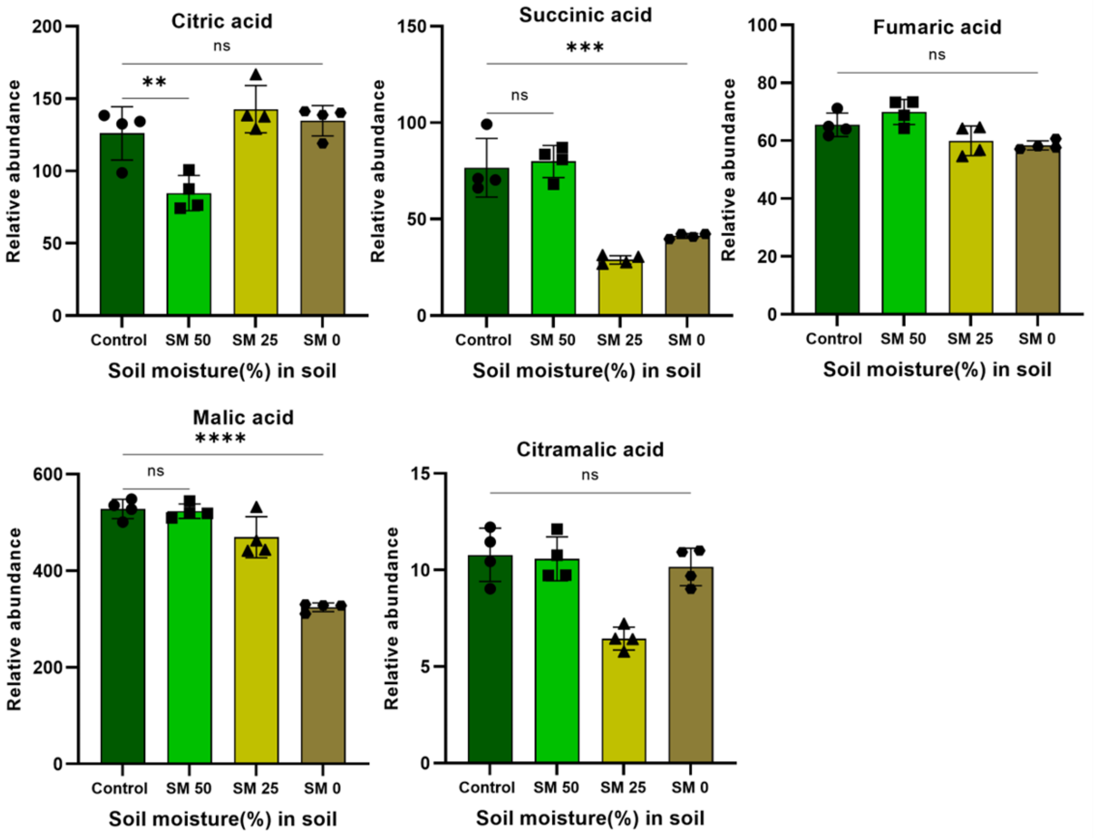


**Supplementary Figure S.2.** Impact of progressive drought on central carbon metabolism and TCA Cycle intermediates in potato leaves. Significant reductions in succinic acid (at SM 25 and SM 0) and malic acid (at SM 0) were observed. Citric acid decreases under mild stress (SM 50) but recovers and increases under more severe stress. Fumaric acid and citramalic acid levels remain stable across all treatments. Data are presented as mean ± standard error mean (n = 4). Significant differences between groups were determined by statistical analysis: ns = not significant, $***p < 0.001$, $****p < 0.0001$

**Supplementary Table 1:** Correlation of Soil moisture value by Lutron PMS-714 with field capacity and interpretation of mean observations of ET satellite values and respective advisories

| Soil moisture Sensor value | Field capacity | Satellite value | Interpretation |
|---|---|---|---|
| ≥ 17 | 100% | ≤0 | Overwatered |
| 15-17 | 80% | 0-1.5 | Watered |
| 14-10 | 50% | 1.5-2.5 | Required water |
| 9-5 | 25% | ≥2.5 | Critical limit |
| 4-0 | 10-5% | | |

**Supplementary Table 2:** Temperature values (Min and Max) in Celsius and average relative humidity values in percentage of field experiment conducted from 15.03.2024-4.04.2025

| Date | Field data | | |
|---|---|---|---|
| | Temperature (°C) | | Relative Humidity(%) |
| | Min | Max | Average |
| **15-03-2024** | 11 | 29.9 | 58.3 |
| **16-03-2024** | 12 | 30 | 60.9 |
| **17-03-2024** | 13 | 30 | 57.7 |
| **18-03-2024** | 12 | 31 | 57.5 |
| **19-03-2024** | 13 | 31 | 57.0 |
| **20-03-2024** | 14 | 32 | 58.1 |
| **21-03-2024** | 17 | 27 | 58.7 |
| **22-03-2024** | 17 | 31 | 69.4 |
| **23-03-2024** | 18 | 33 | 59.3 |
| **24-03-2024** | 16 | 30 | 49.7 |
| **25-03-2024** | 16 | 35 | 57.4 |
| **26-03-2024** | 18 | 32 | 53.0 |
| **27-03-2024** | 20 | 37 | 58.5 |
| **28-03-2024** | 19 | 35 | 70.4 |
| **29-03-2024** | 17 | 35 | 67.9 |
| **30-03-2024** | 14 | 34 | 81.5 |
| **31-03-2024** | 17 | 34 | 70.4 |
| **01-04-2024** | 16 | 31 | 65.9 |
| **02-04-2024** | 15 | 31 | 64.0 |
| **03-04-2024** | 16 | 35 | 54.4 |
| **04-04-2024** | 18 | 36 | 54.5 |
| **05-04-2024** | 9.8 | 36 | 60.2 |

**Supplementary Table 3:** Mean observation reading of soil moisture under control and drought experiment (Lutron PMS-714) and satellite advisories for irrigation (DHI Denmark)

| | Mean observation | | |
|---|---|---|---|
| **Date** | **Control** | **Drought** | **Satellite advisories** |
| **3/22/2024** | 16.7 | 13.4 | Watered |
| **3/23/2024** | 16.7 | 11.6 | Watered |
| **3/24/2024** | 15.0 | 11.0 | Watered |
| **3/25/2024** | 15.0 | 9.7 | Watered |
| **3/26/2024** | 15.0 | 8.4 | Watered |
| **3/27/2024** | 15.1 | 8.4 | Watered |
| **3/28/2024** | 15.8 | 7.2 | Watered |
| **3/29/2024** | 16.8 | 7.1 | Watered |
| **3/30/2024** | 14.3 | 6.6 | Watered |
| **3/31/2024** | 14.5 | 5.4 | Watered |
| **4/1/2024** | 18.2 | 4.0 | Watered |
| **4/2/2024** | 18.4 | 2.6 | Required water |
| **4/3/2024** | 18.2 | 2.0 | Required water |
| **4/4/2024** | 18.0 | 0.5 | Required water |
| **4/5/2024** | 17.3 | 0.0 | Required water |

**Supplementary Table 4:** The metabolite profiles of potato leave under drought stress (SM 50, 25, 0). The metabolite features obtained by MeOX-TMS derivatization of soluble extracts followed by GC-MS spectra. The data represents mean of four replicates. Ribitol is used as an internal standard and set 100 for normalization. SEM represents standard error of mean.

| **S.No.** | **Name** | **RT** | **Control** | | **SM50** | | **SM 25** | | **SM 0** | |
|---|---|---|---|---|---|---|---|---|---|---|
| | | | **Mean** | **SEM** | **Mean** | **SEM** | **Mean** | **SEM** | **Mean** | **SEM** |
| 1 | Lactic Acid | 11.56 | 23.2 | 0.1 | 17.8 | 1.2 | 50.5 | 15.7 | 32.1 | 1.3 |
| 2 | Glycolic acid | 11.86 | 7.9 | 0.5 | 9.5 | 0.5 | 8.1 | 0.6 | 5.0 | 0.2 |
| 3 | Valine | 14.04 | 7.4 | 1.8 | 21.0 | 1.5 | 122.8 | 7.8 | 51.5 | 2.6 |
| 4 | Hydroxylamine | 12.39 | 6.7 | 0.3 | 0.0 | 0.0 | 15.4 | 1.3 | 9.1 | 0.8 |
| 5 | Isoleucine | 15.16 | 3.5 | 2.0 | 0.0 | 0.0 | 44.1 | 2.1 | 77.5 | 3.4 |
| 6 | Ethanolamine | 14.76 | 85.6 | 2.2 | 72.7 | 3.5 | 127.2 | 6.0 | 90.2 | 1.7 |
| 7 | Proline | 13.50 | 0.0 | 0.0 | 121.6 | 16.9 | 679.6 | 32.2 | 19.1 | 0.6 |
| 8 | Glycine | 15.35 | 147.0 | 9.0 | 123.9 | 2.6 | 113.6 | 6.2 | 125.4 | 1.6 |
| 9 | Succinic acid | 15.49 | 98.1 | 9.2 | 79.9 | 4.1 | 28.9 | 1.1 | 41.3 | 0.6 |
| 10 | Glyceric acid | 15.67 | 166.8 | 23.8 | 100.7 | 1.3 | 136.1 | 24.4 | 127.3 | 17.8 |
| 11 | Fumaric acid | 16.00 | 65.5 | 2.0 | 69.9 | 2.2 | 60.0 | 2.6 | 58.4 | 0.8 |
| 12 | Serine | 16.08 | 7.9 | 0.2 | 14.4 | 0.9 | 40.4 | 2.3 | 30.1 | 1.0 |
| 13 | Erythrono 1,4 lactone | 16.32 | 9.8 | 0.4 | 14.8 | 0.8 | 12.4 | 0.5 | 11.7 | 0.6 |
| 14 | Threonine | 16.42 | 9.7 | 1.4 | 6.0 | 0.3 | 23.1 | 2.4 | 26.7 | 0.8 |
| 15 | Erythrose | 17.01 | 17.2 | 6.5 | 0.0 | 0.0 | 102.9 | 5.5 | 50.0 | 2.0 |
| 16 | Citramalic acid | 17.55 | 10.8 | 0.7 | 10.6 | 0.6 | 6.4 | 0.3 | 10.2 | 0.5 |

| | | | | | | | | | | |
|---|---|---|---|---|---|---|---|---|---|---|
| 17 | Malic acid | 17.86 | 527.9 | 10.0 | 523.1 | 7.5 | 469.1 | 21.3 | 324.3 | 4.5 |
| 18 | Erythritol | 17.98 | 62.5 | 5.7 | 89.4 | 7.3 | 34.5 | 0.2 | 26.4 | 0.7 |
| 19 | Aspartic acid | 18.18 | 18.2 | 3.3 | 71.1 | 5.2 | 31.0 | 1.8 | 10.3 | 0.9 |
| 20 | Oxoproline | 18.26 | 34.0 | 4.2 | 34.3 | 1.7 | 116.2 | 5.2 | 94.3 | 1.6 |
| 21 | Aminobutanoic acid | 18.34 | 171.0 | 6.4 | 153.5 | 3.6 | 169.5 | 9.3 | 156.3 | 2.2 |
| 22 | 2,3,4 Trihydroxybutyric acid | 18.42 | 101.4 | 6.6 | 102.0 | 4.2 | 29.6 | 1.1 | 136.3 | 14.4 |
| 23 | Phenylalanine | 19.50 | 81.6 | 19.6 | 71.5 | 3.9 | 119.6 | 6.1 | 58.5 | 1.4 |
| 24 | Ribonic acid | 21.04 | 131.3 | 17.9 | 200.2 | 11.7 | 54.1 | 22.1 | 63.6 | 0.4 |
| 25 | Tartaric acid | 21.46 | 79.9 | 0.5 | 90.3 | 1.6 | 135.0 | 3.5 | 71.5 | 1.2 |
| 26 | Citric acid | 21.89 | 126.0 | 9.2 | 84.7 | 6.1 | 142.8 | 8.2 | 134.8 | 5.3 |
| 27 | Pinitol | 22.01 | 44.3 | 1.3 | 30.6 | 1.5 | 36.5 | 0.6 | 40.6 | 1.7 |
| 28 | Quinic acid | 22.48 | 119.6 | 12.3 | 179.1 | 4.5 | 137.1 | 5.5 | 73.1 | 1.0 |
| 29 | Fructose | 22.77 | 696.9 | 37.2 | 1116.1 | 25.1 | 582.4 | 20.6 | 547.6 | 9.5 |
| 30 | Glucose | 23.61 | 268.3 | 8.1 | 361.3 | 9.9 | 302.2 | 12.8 | 317.9 | 12.0 |
| 31 | Mannitol | 23.81 | 48.6 | 6.5 | 69.7 | 2.6 | 53.9 | 2.1 | 69.6 | 4.3 |
| 32 | Xylose | 24.90 | 177.8 | 4.0 | 225.1 | 16.0 | 178.9 | 18.7 | 167.3 | 10.0 |
| 33 | Galactaric acid | 25.76 | 0.0 | 0.0 | 0.0 | 0.0 | 15.7 | 5.3 | 0.0 | 0.0 |
| 34 | Octopamine | 26.24 | 0.0 | 0.0 | 0.0 | 0.0 | 16.9 | 0.8 | 25.0 | 0.9 |
| 35 | Palmitic Acid | 27.15 | 173.2 | 3.4 | 103.6 | 7.4 | 211.5 | 12.2 | 138.2 | 3.9 |
| 36 | Myoinositol | 28.32 | 294.8 | 3.2 | 329.8 | 5.7 | 575.9 | 29.4 | 337.4 | 7.0 |
| 37 | Tryptophan | 32.47 | 2.9 | 1.0 | 17.8 | 4.1 | 53.1 | 10.4 | 9.8 | 0.3 |
| 38 | Linolenic acid | 33.20 | 30.9 | 1.9 | 0.0 | 0.0 | 24.4 | 8.6 | 0.0 | 0.0 |
| 39 | Stearic acid | 34.11 | 74.3 | 1.9 | 42.3 | 3.9 | 101.3 | 5.8 | 59.6 | 2.2 |
| 40 | Galacturonic acid | 37.55 | 60.9 | 1.6 | 90.2 | 1.6 | 58.3 | 1.8 | 63.4 | 1.4 |
| 41 | Glucopyranoside | 37.75 | 12.6 | 1.3 | 18.6 | 0.6 | 25.3 | 0.7 | 20.4 | 0.6 |
| 42 | Gluconic acid | 38.62 | 31.8 | 31.8 | 0.0 | 0.0 | 44.0 | 1.4 | 19.5 | 1.6 |
| 43 | Mannobiose | 41.09 | 5.9 | 5.9 | 152.2 | 3.7 | 187.6 | 6.2 | 186.4 | 3.0 |
| 44 | Arbutin | 41.78 | 2.8 | 2.8 | 0.0 | 0.0 | 21.7 | 0.3 | 22.7 | 0.4 |
| 45 | Monopalmitin | 41.90 | 56.6 | 1.3 | 33.4 | 1.7 | 80.0 | 2.5 | 61.9 | 1.5 |
| 46 | Sucrose | 42.67 | 55.1 | 34.2 | 453.2 | 4.9 | 215.7 | 4.3 | 212.0 | 2.5 |
| 47 | Maltose | 44.27 | 8.4 | 0.2 | 6.1 | 3.5 | 1.5 | 1.5 | 0.0 | 0.0 |
| 48 | Glycerol monostearate | 45.14 | 48.5 | 1.1 | 15.5 | 3.0 | 56.5 | 2.5 | 36.9 | 1.7 |
